# Type I interferons are important co-stimulatory signals during T cell receptor mediated MAIT cell activation

**DOI:** 10.1101/686170

**Authors:** Rajesh Lamichhane, Henry Galvin, Rachel F Hannaway, Sara M de la Harpe, Fran Munro, Joel DA Tyndall, Andrea J Vernall, John L McCall, Matloob Husain, James E Ussher

**Affiliations:** Department of Microbiology and Immunology, University of Otago, Dunedin, New Zealand; School of Pharmacy, University of Otago, Dunedin, New Zealand; Department of Surgical Sciences, Dunedin School of Medicine, University of Otago, Dunedin, New Zealand

**Keywords:** Mucosal associated invariant T cells, TCR activation, type I interferons, influenza virus, co-stimulation, effector functions

## Abstract

Mucosal associated invariant T (MAIT) cells are abundant unconventional T cells which can be stimulated either via their T cell receptor (TCR) or by innate cytokines. The MAIT cell TCR recognises a pyrimidine ligand, derived from riboflavin synthesising bacteria, bound to MR1. In infection, bacteria not only provide the pyrimidine ligand but also co-stimulatory signals, such as Toll-like receptor agonists, that can modulate TCR-mediated activation. Recently, type I interferons (T1-IFNs) have been identified as contributing to cytokine-mediated MAIT cell activation. However, it is unknown whether T1-IFNs also have a role during TCR-mediated MAIT cell activation. In this study, we investigated the co-stimulatory role of T1-IFNs during TCR-mediated activation of MAIT cells by the MR1 ligand 5-amino-6-D-ribitylaminouracil/methylglyoxal (5-A-RU/MG). We found that T1-IFNs were able to boost interferon-γ and granzyme B production in 5-A-RU/MG-stimulated MAIT cells. Similarly, influenza virus-induced T1-IFNs enhanced TCR-mediated MAIT cell activation. An essential role of T1-IFNs in regulating MAIT cell activation by riboflavin synthesising bacteria was also demonstrated. The co-stimulatory role of T1-IFNs was confirmed using liver-derived MAIT cells. T1-IFNs acted directly on MAIT cells to enhance their response to TCR stimulation. Overall, our findings establish an important immunomodulatory role of T1-IFNs during TCR-mediated MAIT cell activation.

## Introduction

MAIT cells are recently recognised unconventional T cells which express a semi-invariant T cell receptor (TCR) (most commonly Vα7.2-Jα33 in humans and Vα19-Jα33 in mice) and are restricted to antigen presented by MHC class I related protein, MR1 (Huang et al., 2005; Tilloy et al., 1999; Treiner et al., 2003). MR1 is an evolutionally conserved MHC Ib like molecule and presents unstable pyrimidine intermediates derived from 5-amino-6-D-ribitylaminouracil (5-A-RU), a precursor of riboflavin, triggering MAIT cell activation via their TCR (Corbett et al., 2014; Kjer-Nielsen et al., 2012). Consistent with this, MAIT cells have been shown to be activated by a diverse range of riboflavin synthesising bacteria, including *Escherichia coli, Pseudomonas aeruginosa, Klebsiella pneumoniae*, and *Staphylococcus aureus*, as well as some yeasts, including *Candida albicans* and *Saccharomyces cerevisiae* (Dias, Leeansyah, & Sandberg, 2017; Le Bourhis et al., 2013; Le Bourhis et al., 2010). Additionally, MAIT cells express high amounts of the interleukin (IL)-18 receptor and the IL-12 receptor and respond to cytokine signals (IL-12 and IL-18) independent of their TCR (Jo et al., 2014; Le Bourhis et al., 2010; Ussher et al., 2014). Upon activation, MAIT cells produce pro-inflammatory cytokines such as tumour necrosis factor alpha (TNFα), interferon gamma (IFNγ), and IL-17A, as well as cytotoxic molecules (granzymes and perforin) (Billerbeck et al., 2010; Dusseaux et al., 2011; Kurioka et al., 2015; Le Bourhis et al., 2013). The exact effector functions expressed depend upon the mode of activation (Hinks et al., 2018; Lamichhane et al., 2019; Leng et al., 2018). Many *in vivo* studies have established an important role for MAIT cells in protection against bacterial infections (Chua et al., 2012; Georgel, Radosavljevic, Macquin, & Bahram, 2011; Meierovics, Yankelevich, & Cowley, 2013; Wang et al., 2018) and more recently against viruses (Wilgenburg et al., 2018). Therefore, MAIT cells are an important innate-like T cells that bridge the innate and adaptive arms of the immune system.

MAIT cells are abundant in blood and liver in humans but have also been isolated from various mucosal sites, including the intestine (Dusseaux et al., 2011; Leeansyah, Loh, Nixon, & Sandberg, 2014). Since, robust activation can be achieved by both pathogenic and commensal bacteria (Salerno-Goncalves, Rezwan, & Sztein, 2014; Tastan et al., 2018), MAIT cell activation should be tightly controlled to prevent the hyperactivation by commensals and subsequent tissue damage. Indeed, a pure TCR-mediated activation of MAIT cells has been shown to be insufficient to drive sustained activation, with the requirement for additional innate signals (Leng et al., 2018; Slichter et al., 2016; Tang et al., 2013; Turtle et al., 2011). Some recognised innate signals that can modulate activation of MAIT cells via their TCR are IL-7 (Leeansyah et al., 2015; Tang et al., 2013), Toll-like receptor (TLR) agonists (Chen et al., 2017; Ussher et al., 2016), TNFα (Banki et al., 2019), and IL-1β+IL-23 (Tang et al., 2013).

Type I interferons (T1-IFNs) are among the newest and least explored innate signals during MAIT cell activation. It was first shown by van Wilgenburg et al. that adding either IFNα or IFNβ to IL-12 or IL-18 resulted in IFNγ production by MAIT cells (van Wilgenburg et al., 2016). Additionally, T1-IFN signalling was shown to be important for MAIT cell activation during viral infection in both humans and mice (van Wilgenburg et al., 2016; Wilgenburg et al., 2018). However, it is unknown if T1-IFNs also have a co-stimulatory function in TCR-mediated activation of MAIT cells.

In this study, we have shown that the early response of MAIT cells to a TCR signal can be greatly enhanced by T1-IFNs. By using influenza A virus infection as a potent inducer of T1-IFNs and by blocking the T1-IFNs with vaccinia virus receptor B18R, we have confirmed that the overall response of MAIT cells to a TCR signal can be boosted by T1-IFNs produced during viral infections. Furthermore, we demonstrated that T1-IFNs are important during early MAIT cell activation by riboflavin synthesising bacteria. This co-stimulatory effect of T1-IFNs was also evident in liver derived (Ld-) MAIT cells. Further, T1-IFNs had a direct effect on MAIT cells and the co-stimulatory effect was independent of TNFα. Together our findings suggest a significant role for T1-IFNs in driving MAIT cell activation during bacterial, viral, and inflammatory diseases.

## Results

### T1-IFNs are important modulators of MAIT cell activation and function

To test whether T1-IFNs can activate MAIT cells and if they have a co-stimulatory role in TCR mediated MAIT cell activation, we treated human PBMCs with IFNα 2a or IFNβ alone or with the TCR signal, 5-A-RU/MG. Interestingly, treatment with either T1-IFN alone significantly increased the surface expression of CD69, an early activation marker, on MAIT cells compared to the unstimulated control, however, the expression of CD69 in response to either T1-IFN alone was less than to 5-A-RU/MG (Figure 1B); the response to T1-IFNs alone was largely restricted to MAIT cells (S. Figure 1). Both T1-IFNs further enhanced 5-A-RU/MG triggered CD69 expression on MAIT cells, which upon MR1 blocking was significantly reduced to the level of treatment with T1-IFNs alone (Figure 1B). This suggested that the enhanced CD69 expression in response to 5-A-RU/MG plus T1-IFNs was due to the combination of both TCR-dependent and -independent mechanisms. No enhancement was seen in the expression of 4-1BB, another marker of early TCR activation (Wolfl et al., 2007), with either T1-IFN. In contrast, the percentage of MAIT cells expressing 4-1BB was reduced upon treatment with T1-IFNs and 5-A-RU/MG compared to 5-A-RU/MG alone (Figure 1C).

**Figure 1:**
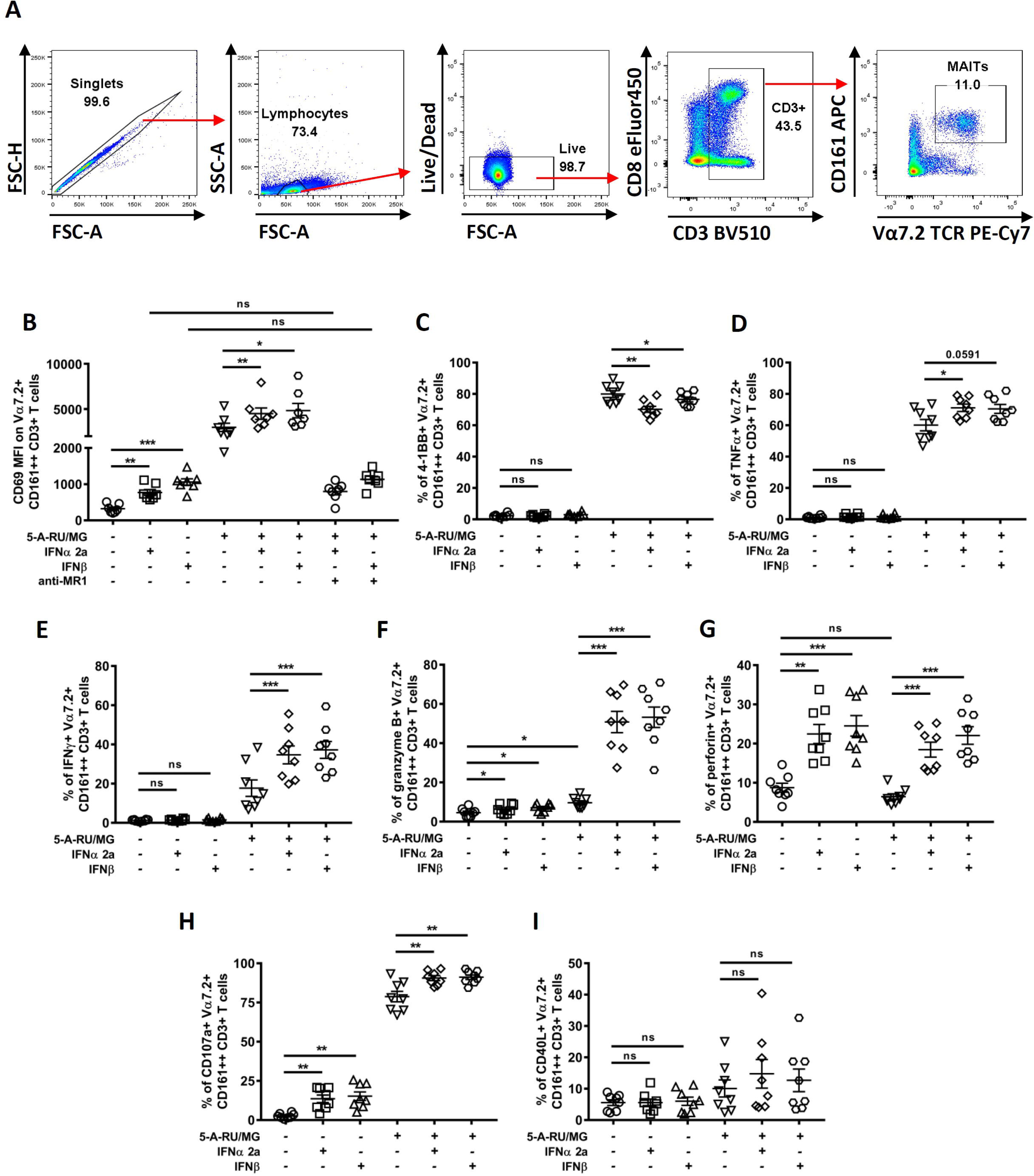
T1-IFNs activate MAIT cells and enhance their response to 5-A-RU/MG. PBMCs were treated with 1nM 5-A-RU/MG or T1-IFNs (450 IU/mL IFNα 2a or 325 IU/mL IFNβ) alone or in combination for 6 hours and CD69 expression (B), and the percentage of MAIT cells expressing 4-1BB (C), TNFα (D), IFNγ (E), granzyme B (F), perforin (G), CD107a (H), and CD40L (I) were assessed by flow cytometry; the gating strategy is also shown (A). MR1 mediated MAIT cell activation was blocked by the addition of anti-MR1 antibody during the treatment (B). Data are presented as mean and S.E.M and are pooled from two independent experiments (n=7 for A, n=8 for B-H). Each data point represents an individual healthy donor. Repeated measures one-way ANOVA with Sidak’s multiple comparison tests were performed for assessing statistical significance. *p<0.05, **p<0.01, ***p<0.001, ns = non-significant.

T1-IFNs in combination with IL-12 or IL-18 have previously been shown to stimulate IFNγ production by MAIT cells (van Wilgenburg et al., 2016). Therefore, we examined whether T1-IFNs can modulate cytokine production by MAIT cells treated with 5-A-RU/MG. Indeed, both IFNα 2a and IFNβ substantially increased the percentage of IFNγ or TNFα producing MAIT cells in response to 5-A-RU/MG (Figure 1D and E). Neither IFNα 2a or IFNβ by itself induced any cytokine production by MAIT cells (Figure 1D and E). This prompted us to investigate if other effector functions can also be boosted by T1-IFNs. In response to 5-A-RU/MG or T1-IFNs alone, more MAIT cells were granzyme B positive than when unstimulated (Figure 1F). Compared to treatment with 5-A-RU/MG alone, the percentage of MAIT cells expressing granzyme B increased by almost four-fold with 5-A-RU/MG + IFNα 2a or 5-A-RU/MG + IFNβ co-treatments (Figure 1F). Interestingly, perforin production in MAIT cells was significantly increased following T1-IFN treatment but not by 5-A-RU/MG (Figure 1G). Adding IFNα 2a or IFNβ to 5-A-RU/MG did not further enhance perforin production compared to treatment with the T1-IFNs alone. T1-IFNs alone significantly induced MAIT cell degranulation, as measured by surface expression of CD107a; compared to treatment with 5-A-RU/MG alone, treatment of MAIT cells with a combination of T1-IFNs and 5-A-RU/MG further increased the proportion of MAIT cells that were CD107a positive (Figure 1H). In contrast, T1-IFNs did not significantly affect CD40L expression, either alone or in combination with 5-A-RU/MG (Figure 1I).

Both CD8^+^ and CD8^−^ MAIT cells responded similarly to T1-IFNs alone or in combination with 5-A-RU/MG, however, CD8^−^ MAIT cells displayed less functional capacity upon activation compared to CD8^+^ MAIT cells, consistent with the recent observation by Dias J et al. (Dias et al., 2018) (S. Figure 2). Taken together, T1-IFNs can directly induce modest MAIT cell activation and augment effector and helper responses of TCR-stimulated MAIT cells.

### Virus modulates the activation and function of TCR-activated MAIT cells

T1-IFNs are produced by infected cells during viral infection (Perry, Chen, Zheng, Tang, & Cheng, 2005; Stetson & Medzhitov, 2006). Therefore, we assessed if virus infection can modulate MAIT cell activation and effector functions. PBMCs were either treated with UV-inactivated IAV alone or in combination with 5-A-RU/MG; uninfected allantoic fluid (AF) was included as a control. IAV alone triggered MAIT cell activation as measured by CD69 surface expression; the activation was specific to IAV treatment as an equivalent volume of AF had no effect on CD69 expression (Figure 2A). Interestingly, adding IAV, but not AF, to 5-A-RU/MG further enhanced CD69 expression on MAIT cells compared to 5-A-RU/MG alone. In PBMCs treated with 5-A-RU/MG + IAV, blocking MR1 significantly reduced CD69 expression, but not completely compared to unstimulated MAIT cells; the level of CD69 expression was similar to the level induced by IAV alone (Figure 2A). IAV alone had little effect on 4-1BB expression on MAIT cells but reduced the 4-1BB expression when added with 5-A-RU/MG compared to 5-A-RU/MG alone (Figure 2B). Neither TNFα nor IFNγ was produced by MAIT cells with IAV alone, however, the percentage of MAIT cells producing IFNγ, but not TNFα, was significantly increased when PBMCs were co-treated with 5-A-RU/MG and IAV compared with 5-A-RU/MG and AF (Figure 2C and D). A small but non-significant increase in granzyme B production was seen with IAV or 5-A-RU/MG stimulation alone. In contrast, IAV together with 5-A-RU/MG triggered robust production of granzyme B in MAIT cells compared to 5-A-RU/MG alone or co-treatment with 5-A-RU/MG + AF (Figure 2E). Perforin production in MAIT cells was increased by IAV alone, but this did not reach significance; co-treatment with IAV and 5-A-RU/MG resulted in a significant increase in perforin expression compared to 5-A-RU/MG alone or co-treatment with 5-A-RU/MG + AF (Figure 2F). MAIT cells also rapidly degranulated in response to IAV; co-treatment with IAV and 5-A-RU/MG resulted in a greater proportion of MAIT cells degranulating than with 5-A-RU/MG alone (Figure 2G). Similarly, co-treatment with IAV and 5-A-RU/MG resulted in a significant increase in CD40L expression by MAIT cells compared to 5-A-RU/MG treatment alone (Figure 2H). Taken together, signals during virus infection induce activation and degranulation of MAIT cells and can further enhance activation, cytokine production, and the cytotoxic potential of MAIT cells stimulated via their TCR, similar to T1-IFNs.

**Figure 2:**
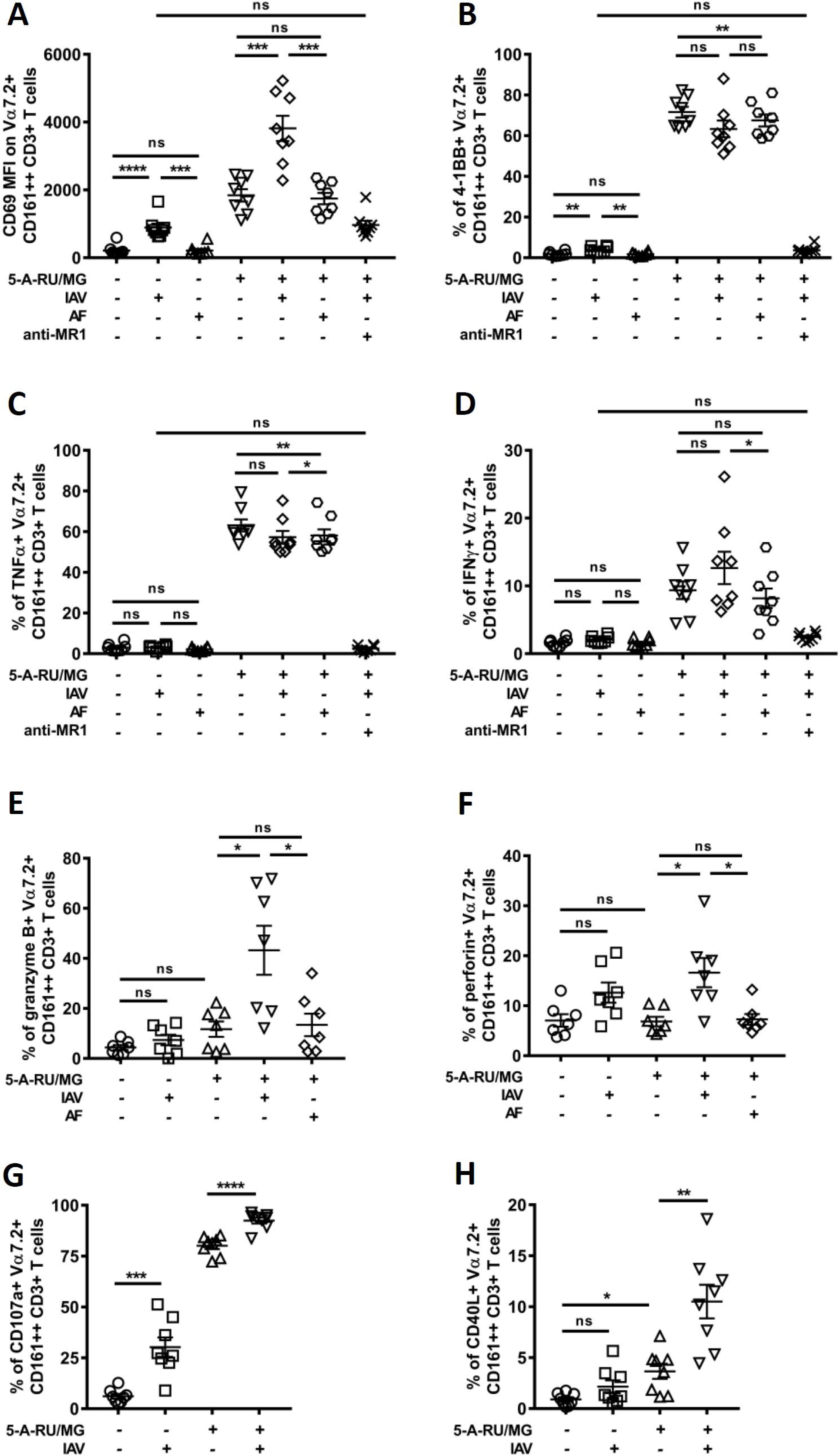
Virus activates MAIT cells and enhances their response to 5-A-RU/MG. PBMCs were treated with 1nM 5-A-RU/MG, UV-irradiated IAV at a MOI of 2, or the equivalent volume of allantoic fluid (AF) alone or in combination for 6 hours and CD69 expression (A), and the percentage of MAIT cells expressing 4-1BB (B), TNFα (C), IFNγ (D), granzyme B (E), perforin (F), CD107a (G), and CD40L (H) were assessed by flow cytometry. In some experiments, MR1-mediated MAIT cell activation was blocked by adding anti-MR1 antibody during the 5-A-RU/MG + IAV co-treatment (A-D). Data are presented as mean and S.E.M and are pooled from two independent experiments (n=8 for A-D, G, and H and n=7 for E-F). Each data point represents an individual healthy donor. Repeated measures one-way ANOVA with Sidak’s multiple comparison tests were performed for assessing statistical significance. *p<0.05, **p<0.01, ***p<0.001, ****p<0.0001, ns = non-significant.

### Virus induced MAIT cell activation, degranulation and hyper-response to TCR signal is dependent upon T1-IFNs

To confirm whether the synergy of 5-A-RU/MG and IAV was mediated via virus induced T1-IFNs, we treated PBMCs with IAV + 5-A-RU/MG and neutralised T1-IFNs with Vaccinia virus protein B18R. Inhibiting T1-IFNs completely abrogated the viral enhancement of CD69, IFNγ, granzyme B, perforin, CD107a, and CD40L expression by TCR-stimulated MAIT cells (Figure 3A, D-H). Furthermore, the reduction of 4-1BB expression on MAIT cells treated with 5-A-RU/MG + IAV compared with 5-A-RU/MG alone was reversed upon B18R addition (Figure 3B). Though no enhancement of TNFα production was seen with IAV and 5-A-RU/MG, blocking T1-IFNs reduced the percentage of MAIT cells producing TNFα (Figure 3C). Furthermore, the increased CD69 surface expression and degranulation observed in MAIT cells when PBMCs were treated with IAV alone were also significantly reduced upon T1-IFN blockade (Figure 3I and J). These findings confirm that the IAV induced MAIT cell activation and enhanced response to TCR stimulation was T1-IFNs mediated.

**Figure 3:**
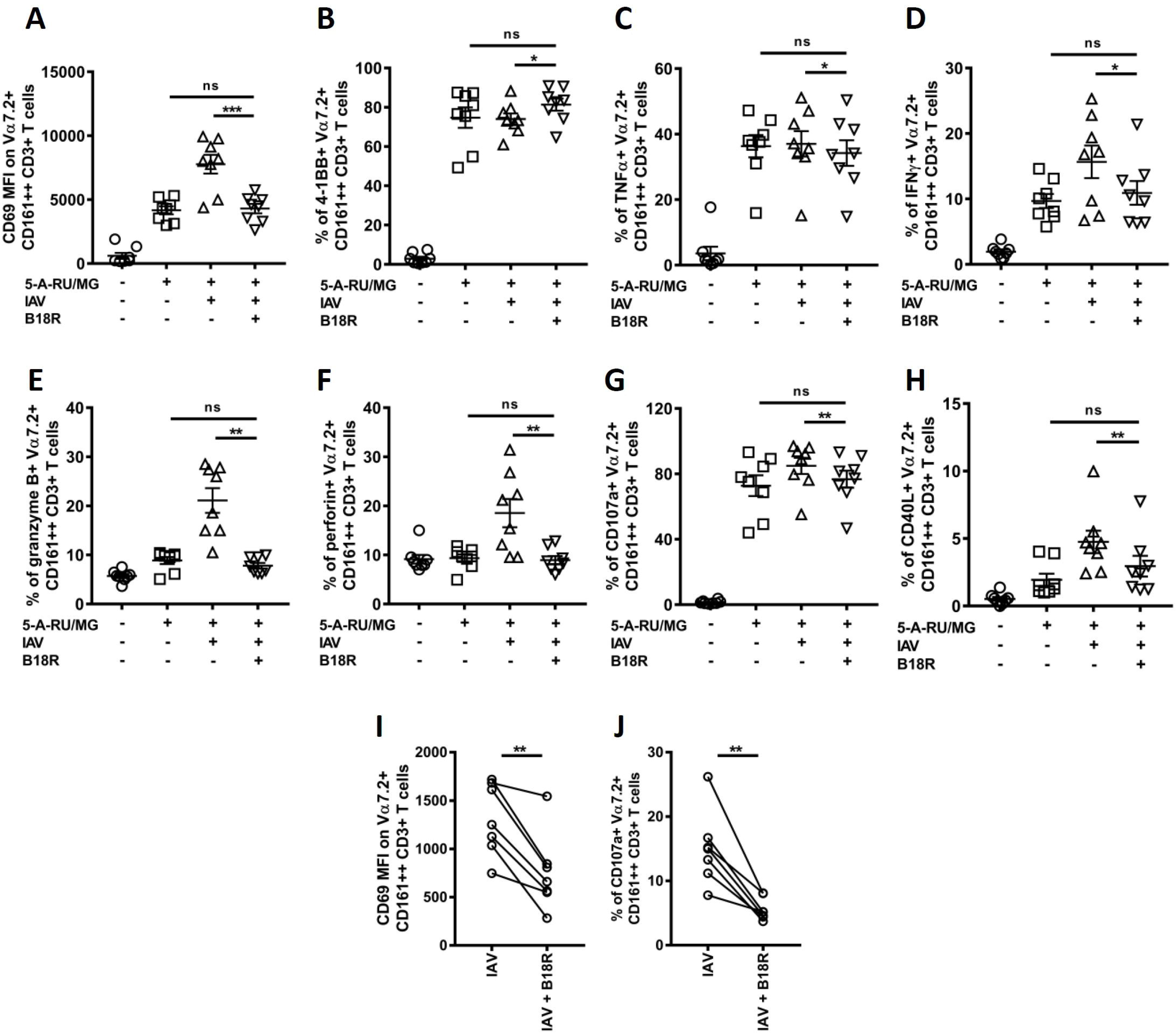
T1-IFNs mediate virus-induced MAIT cell activation and enhancement of the MAIT cell response to 5-A-RU/MG. PBMCs were treated with 1nM 5-A-RU/MG alone or in combination with UV-fixed IAV at a MOI of 2 ± vaccinia virus B18R protein (1 µg/mL) for 6 hours and CD69 expression (A), and the percentage of MAIT cells expressing 4-1BB (B), TNFα (C), IFNγ (D), granzyme B (E), perforin (F), CD107a (G), and CD40L (H) were assessed by flow cytometry. PBMCs were also treated with IAV alone at 2 MOI ± vaccinia virus B18R protein for 6 hours and expression of CD69 (I) and the percentage of MAIT cells expressing CD107a (J) assessed. Data are presented as mean and S.E.M and are pooled from two independent experiments (n=8 for A-H and n=7 for I and J). Each data point represents an individual healthy donor. Repeated measures one-way ANOVA with Sidak’s multiple comparison tests (A-H) or paired two-tailed t-tests (I and J) were performed for assessing statistical significance. *p<0.05, **p<0.01, ***p<0.001, ns = non-significant.

### T1-IFNs are important for early activation of MAIT cells by riboflavin producing bacteria

Bacteria can provide both MR1 ligand and co-stimulatory signals leading to robust activation of MAIT cells (Chen et al., 2017; Ussher et al., 2016). Recently, we reported that various innate signalling pathways, including IFNα/β signalling, were enriched early in MAIT cells upon stimulation with *E. coli* compared to activation with 5-A-RU alone (Lamichhane et al., 2019). Therefore, we activated MAIT cells with *E. coli* for 6 hours with or without B18R to assess the role of T1-IFNs during early MAIT cell activation by a riboflavin synthesising bacteria. Upon activation by *E. coli*, MAIT cells rapidly upregulated surface expression of CD69, 4-1BB, CD40L, and CD107a, produced TNFα and IFNγ, and increased expression of the cytotoxic molecules granzyme B and perforin; all except 4-1BB were reduced upon blocking T1-IFNs, although the change in granzyme B did not reach significance (Figure 4A-H). Interestingly, *E. coli* induced perforin production in MAIT cells was completely inhibited by blocking T1-IFNs (Figure 4F). Taken together, T1-IFNs are important drivers of MAIT cell activation and effector functions during early TCR stimulation by riboflavin synthesising bacteria.

**Figure 4:**
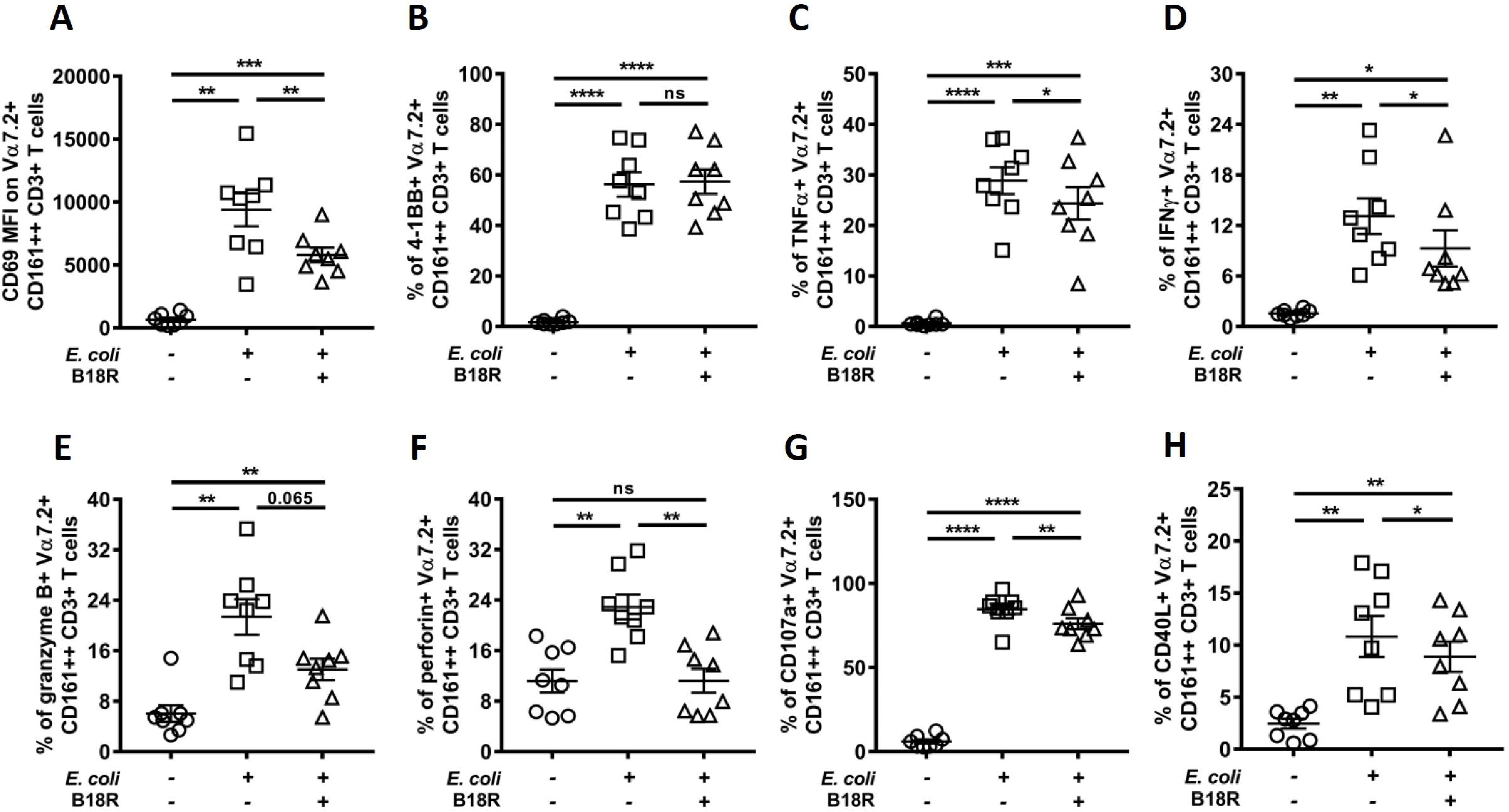
T1-IFN signalling is important for early MAIT cell activation by riboflavin synthesising bacteria. PBMCs were treated with paraformaldehyde fixed *E. coli* at 10 BpC ± 1 µg/mL vaccinia virus B18R protein for 6 hours and CD69 expression (A), and the percentage of MAIT cells expressing 4-1BB (B), TNFα (C), IFNγ (D), granzyme B (E), perforin (F), CD107a (G), and CD40L (H) were assessed by flow cytometry. Data are presented as mean and S.E.M and are pooled from two independent experiments (n=8). Each data point represents an individual healthy donor. Repeated measures one-way ANOVA with Sidak’s multiple comparison tests were performed for assessing statistical significance. *p<0.05, **p<0.01, ***p<0.001, ****p<0.0001, ns = non-significant.

### Liver derived MAIT cells respond to IAV and IFNβ similarly to MAIT cells isolated from blood

Next, to determine whether T1-IFNs are also co-stimulatory signals for liver-derived MAIT cells, we treated liver-derived MAIT cells with IAV, IFNβ or 5-A-RU/MG alone, or in combination (Figure 5); representative flow cytometry plots are shown in S. Figure 2. Liver-derived MAIT cells stimulated with IAV or IFNβ rapidly degranulated and produced more granzyme B and perforin compared to untreated cells, although only CD107a reached significance (Figure 5F-H). The percentages of MAIT cells producing IFNγ and granzyme B were also increased upon co-treatment with IFNβ and 5-A-RU/MG compared with 5-A-RU/MG alone, although only IFNγ reached significance (Figure 5E and F). No significant effect was seen on TNFα and 4-1BB expression following IFNβ or IAV addition to 5-A-RU/MG compared to 5-A-RU/MG alone (Figure 5C and D). Co-treatment with IAV and 5-A-RU/MG also increased CD40L, although this did not reach statistical significance (Figure 5I). Taken together, MAIT cells from both blood and liver can be similarly modulated *in vitro* by T1-IFNs.

**Figure 5:**
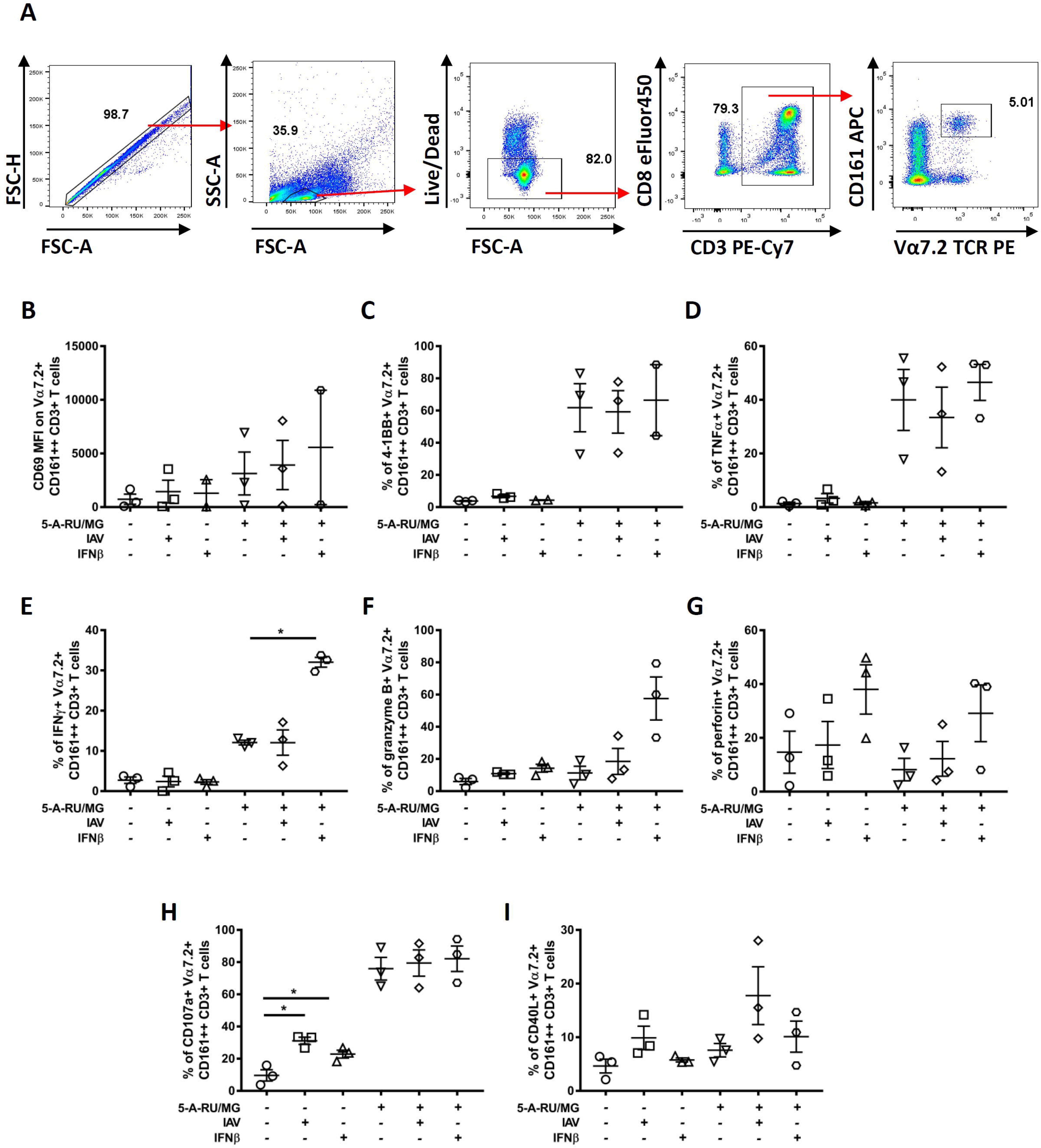
Modulation of liver derived MAIT cell activation by IFNβ and IAV. Human liver derived mononuclear cells were treated with 1nM 5-A-RU/MG or 325 IU/mL IFNβ or IAV at a MOI of 2, alone or in combination, for 6 hours and CD69 expression (B), and the percentage of MAIT cells expressing 4-1BB (C), TNFα (D), IFNγ (E), granzyme B (F), perforin (G), CD107a (H), and CD40L (I) were assessed by flow cytometry; the gating strategy is also shown (A). Data are presented as mean and S.E.M and are pooled from three independent experiments (n=3 except IFNβ ± 5-A-RU/MG treatment in B where n=2). Each data point represents an individual donor. Repeated measures one-way ANOVA with Sidak’s multiple comparison tests were performed for assessing statistical significance. *p<0.05.

### T1-IFN mediated enhancement of TCR stimulation is largely TNFα independent

Monocytes are important mediators of MAIT cell activation by bacterial, viral, or innate signals (Loh et al., 2016; Slichter et al., 2016; Ussher et al., 2016). It is known that monocytes activated with T1-IFNs can trigger more robust MAIT cell activation in response to bacteria (Ussher et al., 2016). To investigate whether T1-IFNs were activating monocytes, we assessed TNFα production by monocytes in PBMCs stimulated with T1-IFNs and/or 5-A-RU/MG. Interestingly, IFNβ, but not IFNα 2a or 5-A-RU/MG alone, triggered significant TNFα production in monocytes (which were mostly CD14^+^HLA-DR^+^) (Figure 6A-C). This was interesting as neither IFNα 2a nor IFNβ alone induced any TNFα production in MAIT cells (Figure 1D).

**Figure 6:**
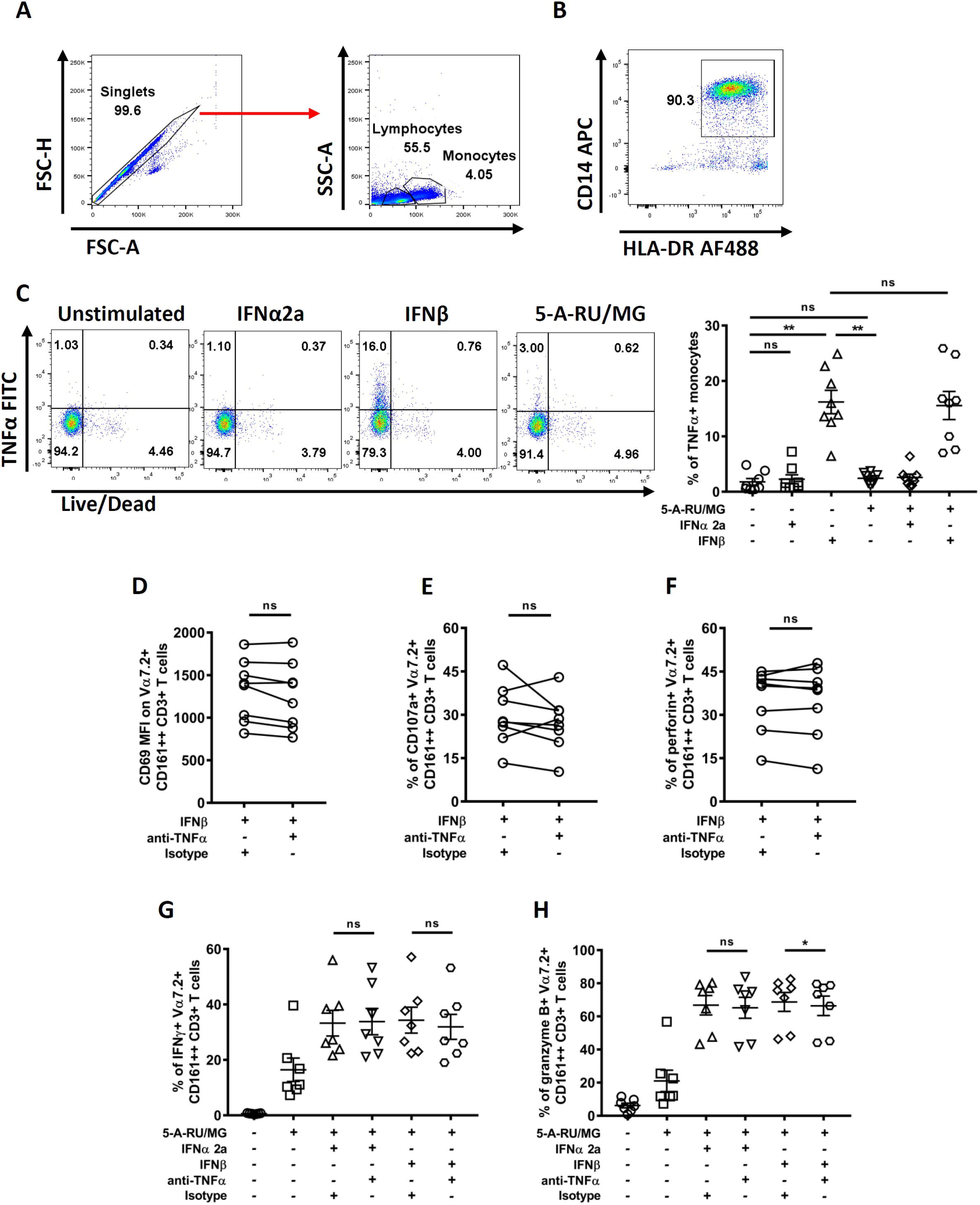
T1-IFN mediated enhancement of the MAIT cell response to 5-A-RU/MG is TNFα independent. PBMCs were treated with 1nM 5-A-RU/MG or T1-IFNs (450 IU/mL IFNα 2a or 325 IU/mL IFNβ) alone or in combination for 6 hours and production of TNFα by monocytes was assessed by flow cytometry. Gating for monocytes is shown (A). Cells in the monocytes gate were predominantly CD14^+^HLA-DR^+^; a representative flow cytometry plot is shown (n=5) (B). Representative flow cytometry plots and cumulative data of TNFα production by monocytes (C). Anti-TNFα antibody (5 µg/mL) or isotype control was added when PBMCs were treated with IFNβ (D-F) or co-treated with 1nM 5-A-RU/MG and either T1-IFN (IFNα 2a or IFNβ) (G and H) for 6 hours. After the treatment, expression of CD69 (D), and the percentage of MAIT cells expressing CD107a (E), perforin (F), IFNγ (G), and granzyme B (H) were assessed by flow cytometry. Data are presented as mean and S.E.M and are pooled from two independent experiments (n=8 for C-F and n=7 for G-H). Each data point represents an individual healthy donor. Repeated measures one-way ANOVA with Sidak’s multiple comparison tests (C, G and H) and two-tailed paired t-tests (D-F) were performed for assessing statistical significance. *p<0.05, **p<0.01, ns = non-significant.

TNFα has been shown to induce upregulation of surface expression of CD69 on MAIT cells (Chiba et al., 2017). Therefore, to test whether the enhanced CD69, CD107a and perforin expression by MAIT cells with IFNβ alone was mediated by TNFα production, we added anti-TNFα antibody with IFNβ treatment. Blocking TNFα had no significant effect on the expression of CD69, CD107a or perforin by MAIT cells (Figure 6D-F). Recently, TNFα produced during treatment of THP1 cells with IgG-opsonized *E. coli* was shown to enhance IFNγ production by MAIT cells (Banki et al., 2019). Therefore, we blocked TNFα in PBMCs co-treated with 5-A-RU/MG + IFNα 2a or 5-A-RU/MG + IFNβ to test whether the synergistic effect seen was mediated by TNFα production. The enhancement of IFNγ production in MAIT cells treated with T1-IFNs was TNFα independent whereas a small but significant reduction was observed in granzyme B production upon TNFα blocking (Figure 6G and H).

### T1-IFNs can directly modulate the response of MAIT cells to TCR stimulation

As the T1-IFN-mediated enhancement of the MAIT cell responses to 5-A-RU/MG was largely TNFα-independent (Figure 6), we hypothesised that T1-IFNs might exert their co-stimulatory effect directly on MAIT cells. To investigate this, we isolated Vα7.2^+^ T cells from human PBMCs and activated them with anti-CD3/CD28 beads alone or in combination with IFNα 2a or IFNβ. MAIT cells produced IFNγ and granzyme B upon activation by anti-CD3/CD28 beads, but not with either T1-IFN alone (Figure 7A-C). Interestingly, both IFNα 2a and IFNβ significantly increased production of IFNγ or granzyme B in CD3/CD28-stimulated MAIT cells compared to CD3/CD28 stimulation alone (Figure 7B). This confirmed that T1-IFNs can directly act on MAIT cells to enhance their response to TCR stimulation.

**Figure 7:**
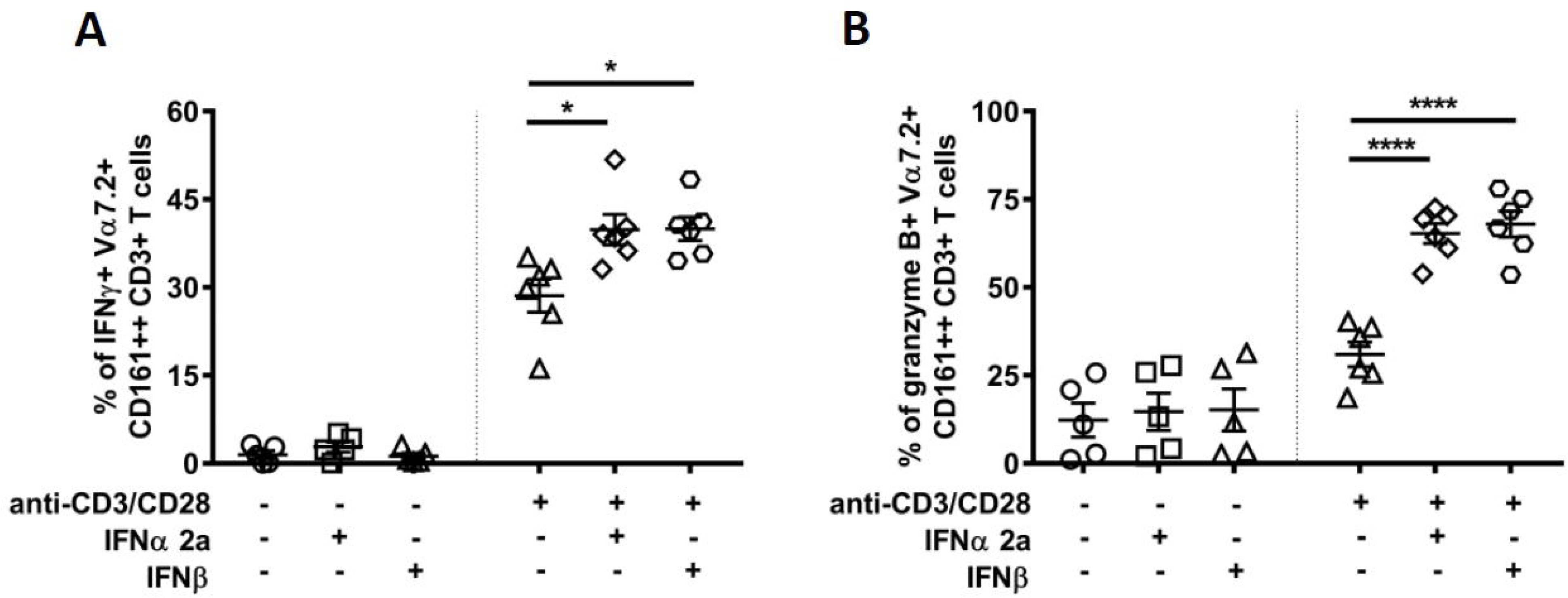
T1-IFNs act directly on MAIT cells. Purified Vα7.2^+^ cells (average purity > 85%) were incubated with anti-CD3/CD28 beads at a ratio of 1:1, or with T1-IFNs (450 IU/mL IFNα 2a or 325 IU/mL IFNβ) alone, or in combination for 6 hours. After treatment, the percentage of MAIT cells producing IFNγ (B), and granzyme B (C) were assessed by flow cytometry; representative FACS plots are also shown (A). Data are presented as mean and S.E.M and are pooled from two independent experiments (n=5-6). Each data point represents an individual healthy donor. Repeated measures one-way ANOVA with Sidak’s multiple comparison tests were performed for assessing statistical significance. *p<0.05, ****p<0.0001.

## Discussion

In this study, we demonstrated that MAIT cells can be activated by T1-IFNs alone, and that T1-IFNs can augment various effector functions of TCR-stimulated MAIT cells. Similar effects on MAIT cell activation were seen with viral stimulation, and this was mediated by T1-IFNs. T1-IFN-mediated enhancement of MAIT cell activation was also seen with bacterial stimulation. A similar role for T1-IFNs was demonstrated with liver-derived MAIT cells. A direct co-stimulatory effect of T1-IFNs on MAIT cells was also established. Together our findings suggest an important role for T1-IFNs in controlling MAIT cell effector responses.

We found that T1-IFNs alone upregulated expression of CD69, granzyme B, and perforin by MAIT cells and induced degranulation, as measured by CD107a expression. No effect on cytokine production or expression of co-stimulatory molecules was seen with T1-IFNs alone. This is consistent with the previous report by Wilgenburg et al., who found that T1-IFNs only induced significant IFNγ production when combined with IL-12 or IL-18 (van Wilgenburg et al., 2016). In another study, various innate cytokines including IFNα were shown to upregulate CD69 expression on MAIT cells (Chiba et al., 2017). Similarly, Spaan et al. demonstrated that adding IFNα or IL-12 to IL-18 significantly increased CD69 expression and IFNγ production by MAIT cells compared to untreated MAIT cells (Spaan et al., 2016). Therefore, T1-IFNs can activate MAIT cells but MAIT cell cytokine production requires stimulation with additional cytokines.

The effect of T1-IFNs on MAIT cell activation has also be shown *in vivo*. In patients with hepatitis C virus (HCV) infection treated with a combination of pegylated IFN and antiviral drugs, higher expression of CD69 was seen on MAIT cells compared to patients receiving antiviral therapy alone (van Wilgenburg et al., 2016). In another study, IFNα-based therapy for chronic HCV infection was shown to increase CD38 expression on MAIT cells (Spaan et al., 2016). The concentration of IFNα in plasma in patients with systemic lupus erythematous positively correlated with CD69 expression on blood MAIT cells (Chiba et al., 2017).

T1-IFNs also modulated MAIT cell effector functions in response to TCR stimulation. Addition of T1-IFNs to 5-A-RU/MG resulted in increased expression of CD69, granzyme B, and perforin, increased degranulation, and enhanced production of IFNγ and TNFα. In contrast, a decrease in 4-1BB expression was seen. T1-IFNs have previously been shown to modulate the function of TCR stimulated T cells. Priming of IGRP-specific CD8^+^ T cells with either IFNα or IFNβ for two hours significantly increased their expression of granzyme B and their ability to lyse target human primary islet cells. (Newby et al., 2017). Similarly, co-treatment with IFNα2 and anti-CD3 antibody enhanced IFNγ production by mouse invariant natural killer T (iNKT) cell hybridomas compared to anti-CD3 antibody alone (Villanueva, Haeryfar, Mallard, Kulkarni, & Sharif, 2015). T1-IFNs also boosted TCR-induced IL-10, but not IFNγ, production in naïve CD4^+^ T cells (Corre et al., 2013).

It has recently been shown that MAIT cells can be activated during viral infection by cytokines, including IL-12, IL-15, IL-18, and T1-IFNs (Loh et al., 2016; van Wilgenburg et al., 2016); this has been discussed in detail in a recent review (Ussher, Willberg, & Klenerman, 2018). In our study, we looked at early events following IAV infection of PBMCs and showed that IAV can weakly activate and induce degranulation of MAIT cells in as little as 6 hours, and that this was mediated by T1-IFNs. In mice, MAIT cells have previously been shown to rapidly upregulate CD69, CD25, and granzyme B during influenza challenge; T1-IFN signalling was shown to be required for the expression of CD25 on MAIT cells (Wilgenburg et al., 2018). In an *in vitro* HCV infection model, neutralizing T1-IFNs with B18R significantly reduced CD69 expression and production of IFNγ and granzyme B by MAIT cells, suggesting T1-IFN co-operate with cytokines such as IL-12 and IL-18 during viral infection (van Wilgenburg et al., 2016). The lack of cytokine production by MAIT cells in response to IAV alone in our experiments likely reflects the earlier time point at which activation was assessed (6 hours vs overnight in the study by van Wilgenburg et al). Collectively, T1-IFNs play an important role in MAIT cell activation during viral infection.

Viral infection enhanced the activation of TCR-stimulated MAIT cells. IAV infection increased the expression of CD69, granzyme B, perforin, and CD40L, degranulation, and production of IFNγ by MAIT cells activated with 5-A-RU/MG. This enhancement of activation was entirely dependent upon T1-IFNs, as demonstrated by blocking T1-IFNs with B18R. This may have implications for MAIT cell activation during lower respiratory tract infections, where co-infection with bacteria and respiratory viruses are common (Kiedrowski & Bomberger, 2018; Lim, Kweon, Kim, Kim, & Lee, 2019) and in chronic hepatitis virus infection, where gut dysfunction can result in microbial translocation and delivery to the liver via the portal circulation (Brenchley & Douek, 2012).

Bacteria derived TLR agonists have been shown to license APCs and result in enhanced MAIT cell activation upon subsequent treatment with *E. coli* (Ussher et al., 2016). Similarly, in a mouse model, co-administration of TLR agonists was required for accumulation of MAIT cells in the lung in response to intra-nasal 5-OP-RU (Chen et al., 2017; Ussher et al., 2016). Interestingly, we recently reported that T1-IFN signalling pathways are upregulated early in MAIT cells activated by *E. coli*, but not by 5-A-RU (Lamichhane et al., 2019). In this study, we demonstrated that T1-IFNs are important early co-stimulatory signals during activation by *E. coli*. A partial reduction in CD69, granzyme B, perforin, and CD40L expression, degranulation, and production of IFNγ and TNFα by *E. coli* stimulated MAIT cells was seen when T1-IFNs were blocked with B18R. Together, this suggests that *E. coli* induce T1-IFNs, which are an important co-stimulatory signal during early TCR-mediated MAIT cell activation. Intriguingly, at this early time point, perforin expression appears to be completely dependent on T1-IFNs. This is consistent with two previous studies that show TCR-independent perforin production by MAIT cells activated by *E. coli* for 6 and 24 hours (Kurioka et al., 2015; Lamichhane et al., 2019).

A similar co-stimulatory role of T1-IFNs was seen with liver derived MAIT cells. Other innate co-stimulatory signals have previously been reported to play a role in the activation of liver derived MAIT cells. Tang et al reported that treating intrasinusoidal and circulating mononuclear cells with IL-7 or a combination of IL-1β+IL-23 boosted the production of TNFα, IFNγ, IL-2, and IL-17A by MAIT cells stimulated with anti CD3/CD28-coupled beads (Tang et al., 2013). In another study, liver derived MAIT cells were the second highest producers of IFNγ, after NK cells, in response to the TLR8 agonist, ssRNA40, with IFNγ production dependent on IL-12 and IL-18 produced by intrahepatic monocytes (Jo et al., 2014). It is of interest that hepatic IL-7 expression in wild-type mice was increased with various TLR agonists (3, 4, 7, and 9), but in comparison was significantly reduced in T1-IFN receptor deficient (IFNAR1^−/−^) mice (Sawa et al., 2009). Consistently, IFNβ has been shown to enhance IL-7 expression in hepatocytes (Sawa et al., 2009). Furthermore, the expression of IFNγ, TNFα, IL-17A, granzyme A/B, and perforin by MAIT cells was increased when PBMCs were treated with IL-7 alone for 48 hours; IL-7 treatment also reversed the MAIT cell effector dysfunction seen in HIV infected patients (Leeansyah et al., 2015). IL-7 can also synergise with IL-12 to enhance granzyme B production in MAIT cells (Wallington, Williams, Staples, & Wilkinson, 2018). Given that IL-7 is induced by T1-IFNs, future studies should investigate whether the co-stimulatory effect observed with T1-IFNs is mediated at least in part by IL-7.

TNFα was considered as a mediator of the co-stimulatory effect observed with T1-IFNs. We have previously reported that pre-treatment of APCs with IFNα increased their ability to stimulate MAIT cells upon subsequent treatment with *E. coli* (Ussher et al., 2016). Monocytes have been shown to produce TNFα when directly activated by IAV-infected epithelium (Loh et al., 2016). A recent study found that IFNα induced TNFα production by monocytes at 24 hours (Provine et al., 2019). TNFα has been shown to directly enhance surface CD69 expression on MAIT cells (Chiba et al., 2017), and to be required for IFNγ production by MAIT cells in response to IL-18 and IFNα or to a replication-incompetent adenoviral vector (Provine et al., 2019). Recently, opsonisation of *E. coli* with IgG was shown to boost IFNγ production by MAIT cells in a TNFα dependent manner (Banki et al., 2019). Here, we demonstrated that IFNβ but not IFNα 2a directly triggered TNFα production by monocytes at 6 hours. However, the enhancement of 5-A-RU/MG-mediated MAIT cell activation by T1-IFNs in our study was not mediated by TNFα, as neutralising TNFα had minimal effect on MAIT cell activation.

The T1-IFN mediated co-stimulatory effect on the TCR-dependent MAIT cell activation observed in this study could occur in two ways. Firstly, T1-IFNs could act on the APCs (including in an autocrine fashion in infection) leading to increased expression of other co-stimulatory molecules and MR1 expression. Indeed, T1-IFN treatment of dendritic cells has been shown to enhance their T cell stimulatory activity through increased expression of CD86 and CD40 and by enhancing MHC-I expression and cross-presentation (Montoya et al., 2002; Spadaro et al., 2012). Alternatively, T1-IFNs could act directly on MAIT cells and modulate their response as has previously been demonstrated with murine CD8^+^ T cells (Le Bon et al., 2006) and IGRP-specific human CD8^+^ T cells (Newby et al., 2017). Using isolated Vα7.2^+^ cells, we showed that T1-IFNs were able to enhance the response of MAIT cells to TCR stimulation with anti-CD3/CD28 beads. Therefore, T1-IFNs can mediate their co-stimulatory effect by acting directly on MAIT cells. However, this does not rule out additional indirect effects of T1-IFNs via activation of APCs or the involvement of other co-stimulatory signals such as IL-7.

In summary, the MAIT cell response to TCR stimulation is restricted and a co-stimulatory signal, such as that provided by T1-IFNs, is essential for full activation. While commensal bacteria may act as a source of MAIT cell TCR ligands, partially activating tissue resident MAIT cells (Leng et al., 2018; Sobkowiak et al., 2019; Tang et al., 2013), pro-inflammatory signals, including T1-IFNs, produced during infection/inflammation may synergise with the TCR signal and tune the overall MAIT cell effector response (Figure 8).

**Figure 8:**
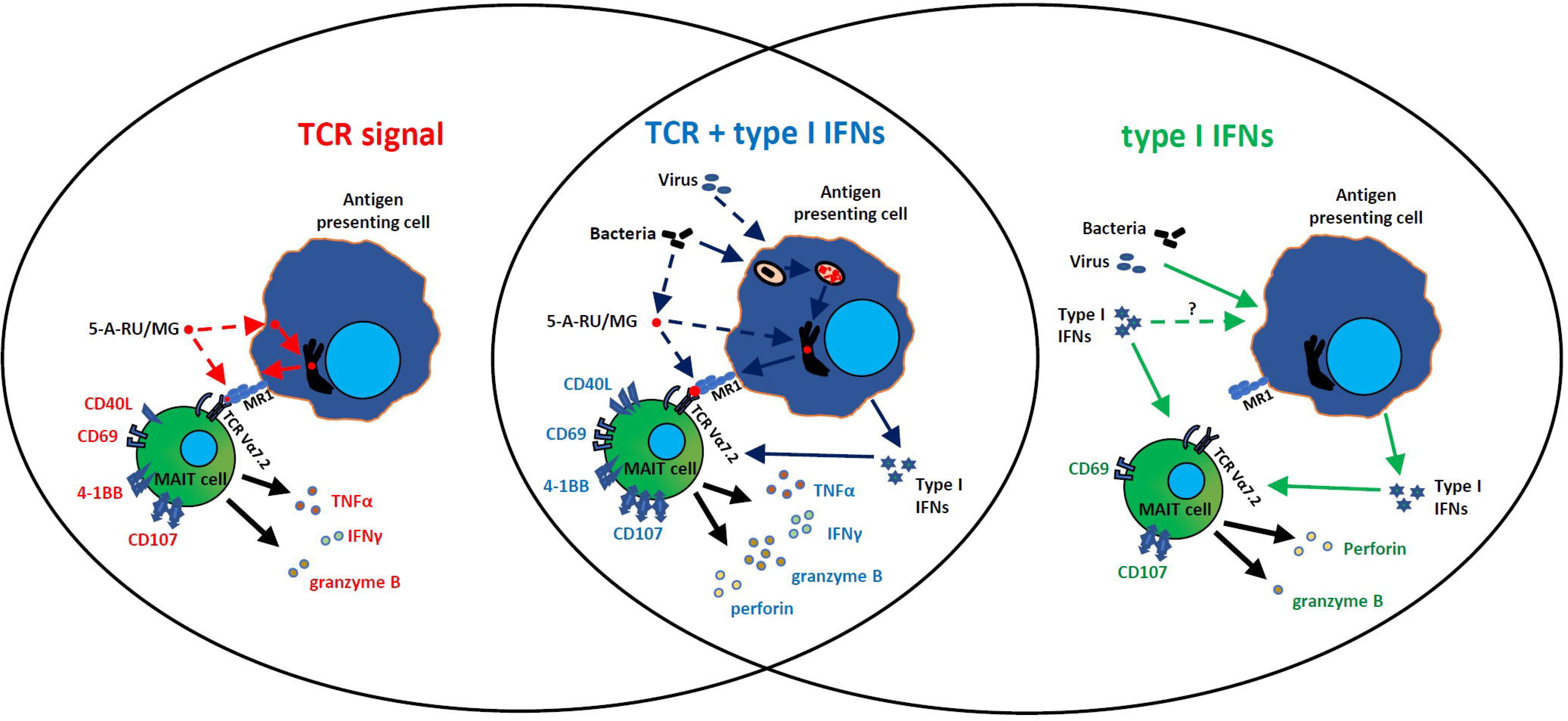
Model of the role of T1-IFNs in MAIT cell activation. The response of MAIT cells to a pure TCR signal (e.g. 5-A-RU/MG), T1-IFNs alone, and the combination of TCR signal and T1-IFNs is shown.

## Materials and Methods

### Mononuclear cell isolation from blood and liver

Peripheral blood was collected from healthy donors and peripheral blood mononuclear cells (PBMCs) were isolated by Lymphoprep™ (Axis-Shield PoC AS, Oslo, Norway) density gradient centrifugation. Isolated PBMCs were stored in liquid nitrogen until use. For each experiment frozen PBMCs were washed and rested overnight in RPMI 1640 pre-supplemented with L-glutamine (Life Technologies Corporation, Carlsbad, USA), 10% fetal calf serum (Gibco, New Zealand), and 100 units/mL Penicillin + 100 µg/mL Streptomycin (Gibco, ThermoFisher Scientific, Waltham, USA). Liver tissue was obtained from two patients with colorectal metastases and one patient with focal nodular hyperplasia undergoing liver resection. A piece of normal liver tissue as far away from the tumour as possible was excised fresh from the main surgical specimen and was cut into smaller pieces, digested with 0.05 mg/mL Liberase TL (Roche, Basel, Switzerland) + 10 units/mL Benzonase nuclease (Merck, Kenilworth, USA) and passed through a 70 µm cell strainer. Liver derived mononuclear cells (LDMCs) were then isolated using Lymphoprep™ density gradient centrifugation. Isolated LDMCs were not cryopreserved but were rested overnight prior to stimulation.

Collection of blood from healthy donors was approved by the University of Otago Human Ethics Committee (H14/046) and collection of liver from patients undergoing hepatic resection was approved by the New Zealand Health and Disability Ethics Committee (16/STH/83). Written informed consent was obtained from each donor.

### Bacteria, virus and MAIT cell ligand

An overnight culture of *Escherichia coli* (*E. coli*) HB101 grown in Luria-Bertani (LB) was fixed with 2% paraformaldehyde for 20 mins at 4°C, washed, and resuspended in phosphate buffered saline (PBS) (Oxoid Ltd, Basingstoke, UK). Fixed bacteria were quantified by flow cytometry using 123 count eBeads (eBioscience, San Diego, USA) on a FACS Canto™ II (BD biosciences, San Jose, USA) and 10 bacteria per cell (BpC) were used for the experiments.

Influenza virus A/PR/8/34 (H1N1) strain (now onwards referred to as IAV) was propagated in 10 day embryonated chicken eggs and titrated on MDCK cells to determine plaque forming units (PFU)/mL (Nagesh, Hussain, Galvin, & Husain, 2017). Aliquots of the viral suspension were stored at −80 °C and irradiated under 30W UV bulb for 30 mins before use in experiments.

MR1 ligand 5-amino-6-D-ribitylaminouracil (5-A-RU) was synthesised as previously reported (Lamichhane et al., 2019). 5-A-RU were stored as single use aliquots at −80 °C. Prior to use, 5-A-RU was mixed with methylglyoxal (MG) at a 1:50 molar ratio (5-A-RU/MG from now onwards) and used immediately.

### MAIT cell activation

A total of 5×10^5^ PBMCs or LDMCs were treated with 1 nM 5-A-RU/MG alone or in combination with either 450 IU/mL human interferon alpha 2a (IFNα 2a) (NR-3083, BEI Resources, Manassas, USA), 325 IU/mL human interferon beta (IFNβ) (NR-3080, BEI Resources) or UV-inactivated IAV (multiplicity of infection of 2) for 6 hours and MAIT cell activation markers (CD69 and 4-1BB) and effector functions (TNFα, IFNγ, CD107a, CD40L, granzyme B, and perforin) were assessed by flow cytometry; IFNα 2a, IFNβ or IAV only treatment or no treatment controls were included as appropriate. For some experiments, PBMCs were treated with *E. coli* (10 BpC) for 6 hours. For antigen presenting cell (APC) free activation, Vα7.2^+^ cells were isolated from PBMCs pre-stained with Vα7.2-PE (3C10, BioLegend) using anti-PE magnetic beads and MS columns (both from Miltenyi Biotech, Bergisch Gladbach, Germany). Isolated Vα7.2^+^ cells were activated with Dynabeads™ human T-activator CD3/CD28 (Gibco, ThermoFisher Scientific) at a ratio of 1:1, with or without IFNα 2a or IFNβ, for 6 hours.

### Blocking antibodies and inhibitors

Anti-MR1 antibody (clone 26.5, BioLegend, San Diego, USA) at 2.5 µg/mL was used to block the MR1-TCR interaction. T1-IFNs were blocked using 1 µg/mL Vaccinia virus B18R recombinant protein (34-8185-81, eBioscience). Brefeldin A (BioLegend) at 5 µg/mL was used to block ER-golgi transport of protein for the assessment of cytokines and cytotoxic molecules. TNFα was blocked with 5 µg/mL anti-TNFα antibody (Mab11, BioLegend); IgG1 isotype control (MOPC-21, BioLegend) was also included.

### Flow cytometry

At the end of the treatment, cells were washed twice with PBS, stained with near IR live/dead stain (Invitrogen, Carlsbad, USA) and antibodies targeting surface markers at 4 ^o^C for 25 mins, followed by two washes with PBS, before fixation with 2% paraformaldehyde at 4 °C for 20 mins. Cells were then washed with PBS and then permeabilization buffer (BioLegend) and stained in permeabilization buffer with antibodies targeting MAIT cell TCR components (Vα7.2 TCR, CD3, and CD8) and intracellular proteins at room temperature for 25 mins. Following additional washes with permeabilization buffer and then PBS, cells were resuspended in PBS. Antibodies used were: anti-CD3 PE-Cy7 (UCHT1, BioLegend), anti-CD3 BV510 (OKT3, BioLegend), anti-CD8 eFluor450 (RPA-18, eBioscience), anti-TCR Vα7.2 PE or PE-Cy7 (3C10, BioLegend), anti-CD161 APC (191B8, Miltenyi Biotec), anti-TNFα FITC (Mab11, BioLegend), anti-IFNγ PerCP-Cy5.5 (4S.B, BioLegend), anti-CD107a PE (H4A3, BioLegend), anti-granzyme B FITC (QA16AO2, BioLegend), anti-perforin PerCP-Cy5.5 (B-D48, BioLegend), anti-CD40L FITC (24-31, BioLegend), anti-4-1BB PE or BV421 (4B4-1, BioLegend), anti-CD14 APC (HCD14, BioLegend), anti-HLA-DR AF488 (L243, BioLegend).

Samples were acquired on a BD FACSCanto™ II or a BD LSRFortessa™ and analysis was performed in FlowJo™ V10 (TreeStar, Ashland, USA). The gating strategy is shown in Figure 1A.

### Statistical analysis

Data were analysed in GraphPad Prism V7.03 and are presented as mean and standard error of mean (SEM). For statistical comparison, paired one-way ANOVA with Sidak’s multiple comparison test between treatments or paired two-tailed t-test between two treatments were used. A two-sided P value of ≤0.05 was considered significant.

## Acknowledgements

This work was supported by grants from the Health Research Council of New Zealand (JEU) and University of Otago Research Grant (JDAT, AJV, JEU). We would like to thank the liver donors and the volunteers who donated the blood samples. The following reagents were obtained through BEI Resources, NIAID, NIH: Human Recombinant Alpha 2a (Alpha A) Interferon (rHuIFN-alpha2a), NR-3083 and Human Interferon Beta (HuIFN-beta), NR-3080.

## Author contributions

JEU and RL designed the project and JEU managed the study. RL, HG, RFH, SMdlH, FM, JDAT, AJV, JM, MH and JEU designed the experiments. RL, HG, RFH and SMdlH performed the experiments. RL and RFH analysed the data. RL and JEU wrote the manuscript. All authors revised and contributed to the editing of the manuscript.

## Conflicts of interest

The authors have no conflicts of interest to declare.

**S. Figure 1: MAIT cells are the predominant CD3^+^ T cell population that are activated by T1-IFNs at 6 hours.** MAIT cells and non-MAIT T cell populations were separated based on Vα7.2 TCR and/or CD161 expression on CD3^+^ lymphocytes (A) and CD69 expression was compared before and after T1-IFN (450 IU/mL IFNα 2a or 325 IU/mL IFNβ) stimulation for 6 hours; representative flow cytometry plot (B) and cumulative data are shown (C). The data presented in Figure 1B for MAIT cells with no treatment or T1-IFN treated (IFNα 2a or IFNβ) are presented here again for comparison with non-MAIT cells. Data are presented as mean and S.E.M and are pooled from two independent experiments (n=7). Each data point represents an individual healthy donor. Repeated measures one-way ANOVA with Sidak’s multiple comparison tests were performed for assessing statistical significance. *p<0.05, **p<0.01, ***p<0.001, ns = non-significant.

**S. Figure 2: Both CD8^+^ and CD8^−^ MAIT cells are modulated similarly by T1-IFNs.** MAIT cells (CD3^+^Vα7.2^+^CD161^++^ T cells) were gated as described in Figure 1A and sub-divided into CD8^+^ and CD8^−^ populations based on CD8 expression; representative flow cytometry plot and cumulative data are shown (A). Percentage of CD8^+^ and CD8^−^ MAIT cells expressing TNFα (B), IFNγ (C), granzyme B (D), and perforin (E) were compared following treatment of PBMCs with 1nM 5-A-RU/MG or T1-IFNs (450 IU/mL IFNα 2a or 325 IU/mL IFNβ) alone or in combination for 6 hours. Data are presented as mean and S.E.M and are pooled from two independent experiments (n=8). Each data point represents an individual healthy donor.

**S. Figure 3: Modulatory role of IFNβ and IAV on liver derived MAIT cell activation.** Liver-derived MAIT cells were identified as CD3^+^ T cells expressing the Vα7.2 TCR and CD161^++^. Representative flow cytometry plots of CD69, 4-1BB, TNFα, IFNγ, granzyme B, perforin, CD107a, and CD40L expression on liver derived MAIT cells following treatment with 1nM 5-A-RU/MG or 325 IU/mL IFNβ or IAV at 2MOI, alone or in combination, for 6 hours are shown.

