## Supplementary figures and images for "Type I interferons are important co-stimulatory signals during T cell receptor mediated MAIT cell activation"

### S. Figure 1

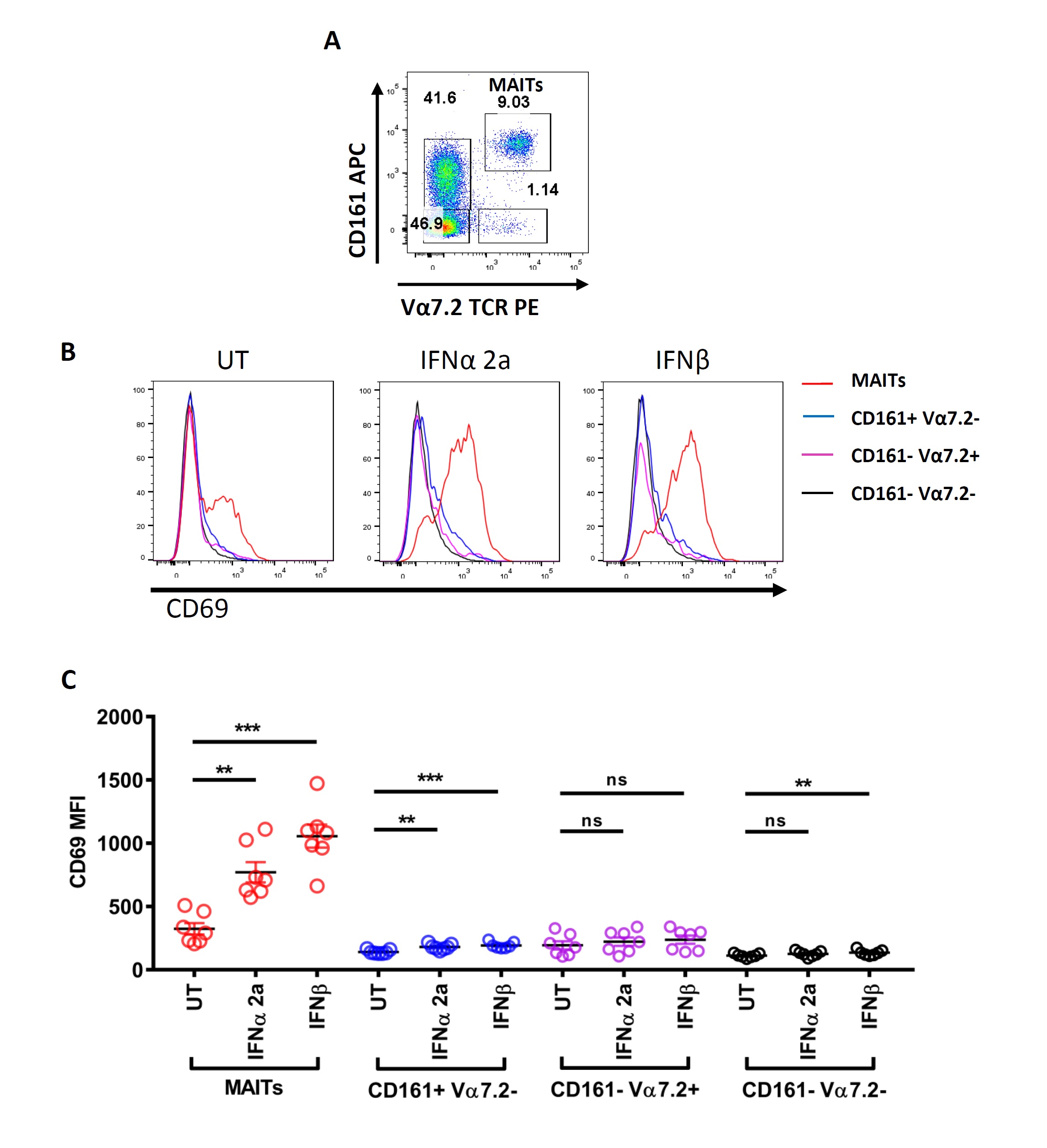

### S. Figure 2

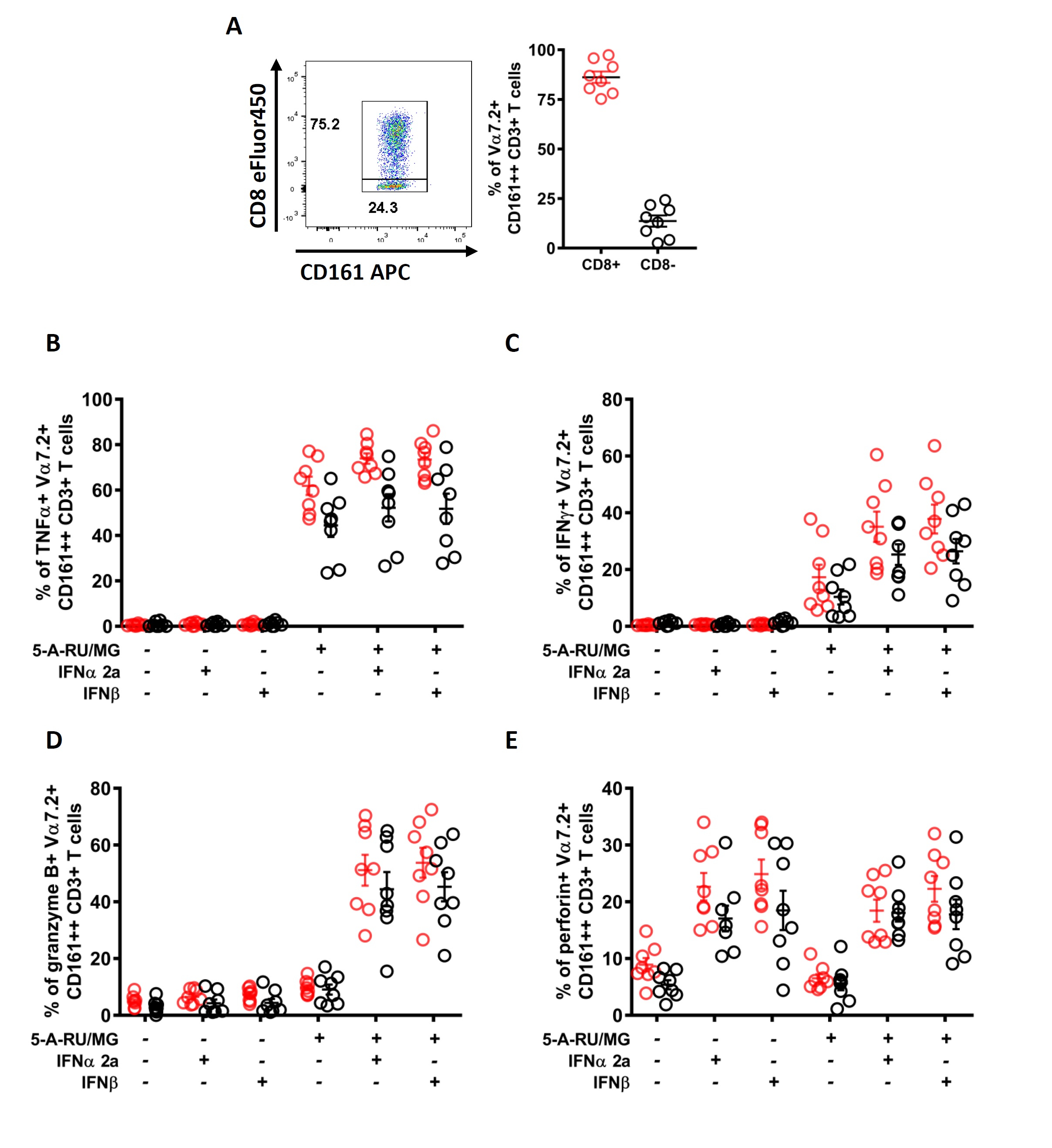

### S. Figure 3

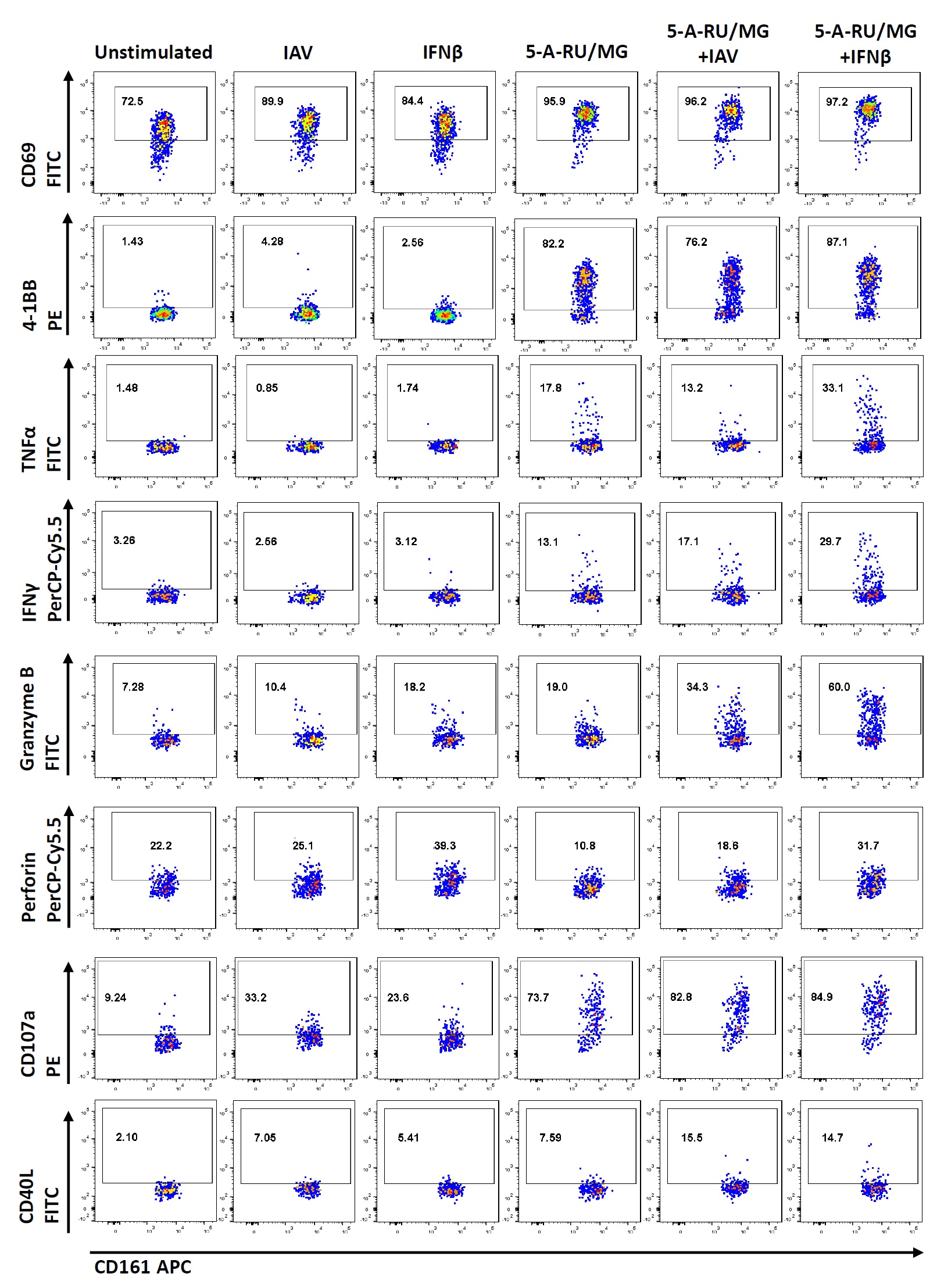
